# Stealth liposomes encapsulating a potent ACAT1/SOAT1 inhibitor F12511: pharmacokinetic, biodistribution and toxicity studies in wild-type mice, and efficacy studies in triple transgenic Alzheimer Disease mice

**DOI:** 10.1101/2023.05.02.539100

**Authors:** Adrianna L. De La Torre, Thao N. Huynh, Catherine C.Y. Chang, Darcy B. Pooler, Dylan Ness, Lionel Lewis, Sanjana Pannem, Yichen Feng, Kimberley S. Samkoe, William F. Hickey, Ta Yuan Chang

## Abstract

Cholesterol is essential to cellular function and is stored as cholesteryl esters (CEs). CEs biosynthesis is responsible by the enzymes acyl-CoA: cholesterol acyltransferase 1 and 2 (ACAT1 and ACAT2), with ACAT1 as the primary isoenzyme in most cells in humans. ACATs are targets for atherosclerosis therapies and may also be promising targets for treating Alzheimer’s Disease (AD). F12511 is a high-affinity ACAT1 inhibitor that has passed phase 1 safety tests for anti-atherosclerosis. Previously, we had developed a nanoparticle system to encapsulate a large concentration of F12511 into a stealth liposome (DSPE-PEG_2000_ with egg phosphatidylcholine). Here, we injected the nanoparticle encapsulated F12511 (nanoparticle F) intravenously (IV) to wild-type (WT) mice and performed HPLC/MS/MS analysis and ACAT enzyme activity measurement. The results demonstrated that F12511 was present within the mouse brain after a single IV but did not over-accumulate in the brain or other tissues after repeated IVs. Histological examination showed that F12511 did not cause overt neurological or systemic toxicity. We then showed that 2-week IV delivery of nanoparticle F to aging 3xTg AD mice ameliorated amyloidopathy, reduced hyperphosphorylated tau and non-phosphorylated tau, and reduced neuroinflammation. This work lays the foundation with nanoparticle F as a possible therapy for AD and other neurodegenerative diseases.

## 1. Introduction

Alzheimer’s disease (AD) is the most common form of dementia. Arguably, the disease has no therapy to slow the clinical progression of the disease. For AD disease modifying therapies, monoclonal antibodies targeting amyloid have advanced to the final clinical trial stage, with aducanumab and lecanemab being the most recently Food and Drug Administration (FDA) approved ^1^. Monoclonal antibodies against tau, as well as against certain inflammatory markers, are also in the clinical pipeline. Besides antibodies, the conventional use of small molecules comes with challenges in crossing the blood brain barrier (BBB) (∼98% of small molecules are not able to cross the BBB) ^2^, as well as possible off-target effects and toxicity. However, one of the key benefits of small molecules is their relatively simple manufacturing and cost-effectiveness when compared to biologics. With a disease as complex as AD, it is likely that a cocktail of therapies will be needed to slow the progression and ultimately cure the disease.

AD is classified as early-onset (EOAD) or late-onset (LOAD), with 99% being LOAD. AD pathological hallmarks consist of extracellular amyloid plaques composed of amyloid beta peptides, neurofibrillary tangles composed of misfolded and hyperphosphorylated tau. Lipid granules also accumulate within the glia, as reviewed in ^3^. LOAD involves multiple genetic risk factors, among them, *APOE* ε4, *CLU*, *ABCA7*, and *ABCA1* are all involved in lipid metabolism ^4, 5–7^. Thus, AD can be considered as a special lipid disease.

Cellular cholesterol homeostasis is under tight control mechanisms involved in uptake of exogenous cholesterol-rich substances, biosynthesis of endogenous cholesterol, intracellular transport, and efflux of cholesterol; the excess cellular cholesterol is stored as cholesterol esters (CEs). (For reviews see ^8–11^. Normally, CE levels in the brains are very low. However, in the vulnerable regions of brain samples with LOAD, CE levels arose by 1.8-fold ^12^. Similar results were observed in three EOAD mouse models ^12, 13^. Additional studies show that in AD patient-derived neurons, increase in CE contents correlates positively with tau pathology ^14^. CEs are biosynthesized by the enzymes acyl-coenzyme A: cholesterol acyltransferases [abbreviated as ACATs] (also named as sterol O-acyltransferases [SOATs] in GenBank. In Genbank ACATs are assigned to acetyl-CoA acetyltransferases, which are separate enzymes that produces acetyl CoA, but not CEs.). There are two ACAT/SOAT genes, encoding two homologous enzymes ^15–18^. Both enzymes use long-chain fatty acyl-CoAs and sterols as substrates ^19^. ACAT1 is expressed in essentially all cells, including cells in the brain; ACAT2 is mainly expressed in intestinal enterocytes and hepatocytes. In various cell types in the CNS, the gene expression levels of ACAT1 are much higher than those of ACAT2, (data retrieved from public domain can be found in ^20^. ACAT1 and ACAT2 are membrane proteins located in the endoplasmic reticulum (ER) and both enzymes are allosterically activated by sterols ^21–24^. In addition, in microglia isolated from various neurodegenerative diseases, and in vulnerable regions of human brains from LOAD, the *Acat1/Soat1* gene is modestly induced (data retrieved from public domain can be found in ^20^.

At the preclinical level, evidence from several laboratories implicates ACAT1 as an important molecular target for the treatment of AD ^25–32^. Mechanistically, ACAT1 blockade by small molecule inhibition or by genetic inactivation (A1B) was reported to offer multiple benefits to AD models including the following: (**1**). A1B increased the content of the neuroprotective oxysterol 24(S)-hydroxycholesterol in the AD mouse brain ^27^ and in the AD patient iPSC derived human neurons ^30^. (**2**). A1B increased autophagy flux and led to the clearance of Aβ oligomers in microglia ^29^; it also increased the clearance of misfolded tau in neurons ^32, 33^. (**3**). In EOAD patient-derived neurons, A1B reduced CE content and prevented their inhibitory effects on tau proteostasis ^30^. (**4**). A1B decreases the protein content of mutant full-length hAPP in the brain of an EOAD mouse model ^27^, and in AD patient iPSC derived human neurons ^30^. (**5**). A1B cleared CEs accumulated in myelin debris treated microglia that lack TREM-2, which is a risk factor for LOAD ^31^. Genetic variants of human *Acat1* have been studied by several research groups. To cite one example, among the four common *Acat1* single-nucleotide polymorphism (SNP) investigated, one exhibits protective haplotype, and one exhibits risk haplotype for dementia development ^34^. In addition, at least one human who has putative homozygous knockout mutation for *SOAT1 ha*d been identified without obviously noticeable issues ^35^, supporting the idea that A1B may not cause overt toxicities in humans. Regarding small molecule ACAT inhibitors: ACAT is a drug target to treat atherosclerosis and many ACAT inhibitors were produced as anti-atherosclerosis agents. Three of these inhibitors, CI1011, K604, and F12511, had passed clinical phase 1 safety tests ^36^. They were all abandoned, some of them (CI1011) for their lack of efficacy to act as a supplement to statin to further reduce serum cholesterol levels ^37^; F12511 or K604 were also withdrawn from clinical trials without providing reasons. **CI1011** is a weak ACAT inhibitor (Ki = 20 µM for both ACAT1 and ACAT2) ^38^. **K604** is a high-affinity ACAT1-specific inhibitor with Ki = 0.45 µM ^38^. **F12511** is a high-affinity inhibitor of ACAT1 (Ki = 0.039 µM); it also inhibits ACAT2 but with less potency (Ki = 0.110 µM) ^39^. **CP113,818** is a high-affinity ACAT inhibitor ^40^(Ki = 0.02 µM for both ACAT1 and ACAT2 ^39^. When CP113,818 was delivered by implanting as part of a custom designed slow-releasing pellet underneath the mouse skin, it was effective in suppressing amyloidopathy in a mouse model for EOAD ^26^, This result strongly suggesting that ACAT inhibitors can be effective to treat AD. Unfortunately, to our knowledge, composition of the slow releasing pellet remained as a company proprietary information. In addition, as an anti-atherosclerosis drug candidate, CP113,818 had failed at the pre-clinical, because it accumulated in adrenals in animals and caused toxicity. It was not clear what causes CP113,818 to be toxic. CP113,818 possesses an asymmetric carbon; this asymmetry is needed for CP113,818 to act as an ACAT inhibitor. The enantiomeric isomer of CP113,818 (and its closely related structural analogs) is inactive as ACAT inhibitor, but also causes severe adrenal toxicity ^41, 42^, suggesting that the toxicity of CP113,818 and its closely structurally analogs such as ATR-101 ^43, 44^ was mainly due to their physicochemical characteristics unrelated to their ability to inhibit ACAT ^41, 42^. Supporting this interpretation is the evidence that the adrenal functions of mice genetically knocked out of ACAT1 is normal ^45^. To overcome adrenal toxicity by CP113,818, Pierre Fabre pharmaceutical produced F12511 (abbreviated as “F”). F12511 also contains an asymmetric center (the chemical structures of CP 113,818 vs F12511 can be found in ^46^; and is a high-affinity inhibitor of ACAT1 (Ki = 0.039 µM). Unlike CP113,818, F12511 passed phase 1 safety tests for anti-atherosclerosis compounds ^36^.

The other high-affinity ACAT1 inhibitor, K604 (the chemical structure of K604 can be found in ^46^), has also passed phase 1 safety tests for anti-atherosclerosis in humans. K604 is known to be impermeable to the mouse brain ^47^. However, whether F12511 is permeable to the brain was unknown. Here, we encapsulated F12511 or K604 in the DSPE-PEG_2000_/PC mixed liposome ^48^ and delivered each to wild type mice by IV injections and performed HPLC/MS/MS analysis and ACAT enzyme activity measurement. We also performed 2-week IV delivery of nanoparticle F to aging 3xTg AD mice to determine if nanoparticle F can ameliorate amyloidopathy, tauopathy, and neuroinflammation, the three major biomarkers for LOAD. The results are reported in the current manuscript.

## 2. Materials and Methods

### 2.1 Ethical handling of animals

The Institutional Animal Care and Use Committee (IACUC) at Dartmouth approved all mouse experiments under the protocol (#00002020). Genetically ablated *Acat1 −/−* mice ^49^ are on C57BL/6 genetic background. Triple transgenic Alzheimer’s disease (3XTg AD) mice ^50^ with or without *Acat1* ^27, 28^ maintained in the Chang lab are on a mixed 129:C57BL/6 genetic background. Mice were subjected to treatment by tail-vein intravenous injection or retro-orbital at 10mL/kg.

### 2.2 Lipids, ACAT inhibitors, solvents, and chemicals

DSPE-PEG_2000_ was from Laysan Bio, Inc (mPEG-DSPE, molecular weight 2,000). L-α-Phosphatidylcholine (from egg yolk) was from Sigma-Aldrich (Catalog No. P-2772). F12511 and K604 were custom synthesized by WuXi AppTec in China. Based on HPLC/MS and NMR profiles, F12511 was 98% in chemical purity and in stereospecificity (F12511 contains an asymmetric center) ^51^; K604 was 98% in chemical purity. CP113,818 was a research gift from Pfizer. [1,1-dioctadecyl-3,3,3,3-tetramethylindotricarbocyanine iodide] DiR (catalog #22070) was purchased from AAT Bioquest.

### 2.3 Nanoparticle formation

Protocol was described in detail in ^48^. Briefly, DSPE-PEG_2000_ dissolved in ethanol (EtOH) and phosphatidylcholine (PC) dissolved in chloroform were combined under vortex. ACAT inhibitors (F12511 or K604) or DiR tracer were dissolved in EtOH were then added to the DSPE-PEG_2000_/PC mixture while under rapid mixing. The final solution contained 30 mM DSPE-PEG_2000_, 0-6 mM PC, and 6-12 mM ACAT inhibitor or 8.3 mM DiR. The final solution was lyophilized overnight and stored away from light at −20 °C. Prior to use, the frozen sample was re-suspended in sterile 1x phosphate-buffered saline (PBS), and bath sonicated in a Branson, 2510 sonicator at 4 °C. The sonicated solution kept in sterile condition, was collected in sterile Eppendorf tubes, and centrifuged at 12,000 rpm for 5 minutes to remove unincorporated materials. The supernatant contained the sterile nanoparticles that were used for treatment directly. Initial studies using DSPE-PEG_2000_ nanoparticles with F12511 at 4-6 mol% without PC followed the same lyophilization procedure; however, after re-solubilization in 1mL PBS, nanoparticles were probe-sonicated on ice under sterile condition, using a Branson probe sonicator, for 2 times 1 min pulses with 5 min rest between sets. The nanoparticles were then used for injection.

### 2.4 Mixed liposomal ACAT activity assay In vitro

This method was as described in ^52^; its use in mouse tissues was described in ^27, 28^. Mouse tissues were homogenized using the Next Advance bullet blender homogenizer with stainless steel beads. The buffer used contained 2.5% 3-((3-Cholamidopropyl) dimethylammonio)-1-propanesulfonate (CHAPS), 1M KCl in 50 mM Tris at pH 7.8. Aliquots of tissue homogenates were then transferred to prechilled glass tubes that contained mixed liposomal mixture of taurocholate, phosphatidylcholine (egg yolk), and cholesterol. Tubes were vortexed and kept on ice. Samples were then incubated in a 37 °C shaking water bath with ^3^H-oleoyl CoA added to start the enzyme reaction for 10 minutes. The assay was stopped by adding CHCl_3_: MeOH (2:1), vortexed, and centrifuged at 500 rpm for 10 minutes. The top phase was removed, and the bottom phase was blown dried by N_2_, 100µl of ethyl acetate was added to each tube, vortexed and spotted on a thin layer chromatography (TLC) plate (Miles Scientific Silica gel HL, Catalog No. P46911). The TLC solvent system used was petroleum ether:ethyl ether:acetic acid (90:10:1). The cholesteryl ester band with R_f_ value at 0.9 was visualized by iodine staining, scraped from the plate, and measured by scintillation counter for radioactivity.

### 2.5 Histology

Wildtype C57/BL6 mice were either left untreated, treated with control DSPE-PEG_2000_/PC nanoparticles, or nanoparticles that contain F12511 (Nanoparticle F). Control and Nanoparticle F treated mice received 7 daily IV injections and were sacrificed 48 h after the last injection. Mice were anesthetized with Avertin (2% in PBS) and, after confirming by lack of toe pinch reflex, were slowly perfused with 10 mL of 4% sucrose in PBS followed by 10 mL of 4% paraformaldehyde (PFA) in 4% sucrose solution in PBS. Tissues were collected and kept in PFA at 4°C on a sample mini rotator overnight before switching solution to 4% sucrose in PBS. Tissues were then collected into histology cassettes, embedded into paraffin blocks, and sectioned onto slides with hematoxylin and eosin (H&E) stain.

### 2.6 HPLC/MS/MS

F12511 was quantified in mouse tissues via LC-MS/MS, CP113,818 (a research gift from Pfizer), a closely related ACAT inhibitor was used as internal standard. Both compounds were dissolved in DMSO at 10 mg/mL and stored at −40 ⁰C. Subsequent working dilutions were made in acetonitrile (ACN) fresh daily. Calibrators and quality controls were made in the appropriate matrix: C57/BL6 plasma (anticoagulant: K3-EDTA, Innovative Research), brain homogenate, liver homogenate, adrenal gland homogenate. Tissues were homogenized at 0.1 g/mL in diH_2_O using stainless steel beads and a Next Advance Bullet Blender. All samples (50µL) were protein precipitated, with 150 µL of 10 ng/mL CP113,818 in ACN as internal standard added via vortex 1 min and centrifugation 5 min at 15,000 rpm. 150 µL of supernatant was transferred to amber autosampler vials and 10 µL was injected on the LC-MS/MS system. HPLC separation was achieved with isocratic conditions of 5% diH_2_O, 95% methanol, and 0.1% formic acid over 2.5 min at a flow rate of 1.5 mL/min on a Phenomenex Luna C18 100 x 4.6mm, 3 micron column fitted with a 10 x 4 mm C18 guard at 40⁰C. A TSQ Vantage mass spectrometer was operated in positive ion mode with a collision pressure of 1.3 mTorr to measure F12511 (470.242→268.120 m/z) and CP113,818 (471.177→201.040 m/z) with collision energies of 16 and 21, and SLens values of 139 and 187, respectively. The heated ESI source was operated with a spray voltage of 4500 V, vaporizer temperature at 500 ⁰C, capillary temperature at 250 ⁰C, sheath and auxiliary gases at 30 and 15 arbitrary units, respectively. The quantitative range for F12511 was 0.3-1000 ng/mL for samples from liver, adrenal glands, brain, and whole blood, and 0.5-1000 ng/mL for plasma.

### 2.7 Preparation of brain homogenates and western blot analysis

Half of forebrains from male and female 3xTg AD mice at the indicated age ranges were homogenized at 4 °C in the Bullet Blender with stainless steel beads in sucrose buffer with protease inhibitors as described in previously published protocol ^53^. Samples were centrifuged at 4 °C for 5min at 12,000 rpm to collect any precipitate. Homogenates were then collected for protein analysis by Lowry assay and divided into pre-chilled Eppendorf tubes and stored at −80 °C until ready for analysis. For Western blot analysis, loading dye was added such that the final sample loading concentration contained at least 10% sodium dodecyl sulfate (SDS). Western blots for detecting human mutant full-length human mutant APP were described in ^27^; Western blots to detect human tau and hyperphosphorylated tau were described in ^32^. Blots were developed using Odyssey DLx LI-COR and analyzed using Image Studio Software by LI-COR.

### 2.8 Antibodies

Mouse anti-human amyloid beta_1-42_ (6E10), mouse anti-human tau (HT7) and mouse anti-human hyperphosphorylated tau [PHF-tau (AT8)], and anti-beta-actin (beta-actin as loading control) were from Covance, Thermo Fisher, and Sigma, respectively.

### 2.9 *ELISA* for monitoring huma amyloid beta _1-42_ (Aβ1-42)

Tissues homogenates were prepared as described above. Homogenates underwent formic acid extraction following previously published protocol ^54^. Samples were then measured by using human Aβ1-42 enzyme-linked immunoassay (ELISA) kit (Catalog No. KHB3441, Invitrogen by Thermo Fisher Scientific).

### 2.10 Monitor biodisitrbution of nanoparticle with DiR

Method was adapted from ^55^ with modifications. Briefly, 3 months old wild-type female mice were IV injected with DiR nanoparticle. 4 h later, mice were sacrificed and perfused with ice cold 1xPBS. Brains were then collected and separated into forebrain and cerebellum regions. Each brain region was then homogenized in 5mM EDTA, 5% Acetone at 4 °C in the Bullet Blender with stainless steel beads and loaded into capillary tubes microhematocrit non-heparinized (Fisher Scientific, Hampton, NH) for fluorescence quantification. Images were captured on the Pearl 500 imaging system (LICOR Biosciences, Lincoln, NE) and quantified using Fiji-ImageJ Software.

### 2.11 Brain Luminex Cytokine Analysis

Cytokines from mouse brain homogenates prepared in section 2.7 were measured using Millipore mouse cytokine multiplex kits (EMD Millipore Corporation, Billerica, MA). Calibration curves from recombinant cytokine standards were prepared with threefold dilution steps in the same matrix as the samples. High and low spikes (supernatants from stimulated mouse PBMCs and dendritic cells) were included to determine cytokine recovery. Standards and spikes were measured in triplicate, samples were measured once, and blank values were subtracted from all readings. All assays were carried out directly in a 96-well filtration plate (Millipore, Billerica, MA) at room temperature and protected from light. Briefly, wells were pre-wet with 100 µl PBS containing 1% BSA, then beads together with a standard, sample, spikes, or blank were added in a final volume of 100 µl and incubated together at room temperature for 30 min with continuous shaking. Beads were washed three times with 100 µl PBS containing 1% BSA and 0.05% Tween 20. A cocktail of biotinylated antibodies (50 µl/well) was added to beads for a further 30 min incubation with continuous shaking. Beads were washed three times, then streptavidin-PE was added for 10 min. Beads were again washed three times and resuspended in 125 µl of PBS containing 1% BSA and 0.05% Tween 20. The fluorescence intensity of the beads was measured in using the Bio-Plex array reader. Bio-Plex Manager software with five-parametric-curve fitting was used for data analysis.

## 3. Results

### 3.1 F12511 is present in the brain after single IV injection of nanoparticle F

We encapsulated F12511 or K604 as part of a DSPE-PEG_2000-_based nanoliposome (also called a “stealth liposome”; when delivered to the blood, DSPE-PEG_2000_ protects the liposome from rapid hepatic degradation *in vivo*). Nanoliposomes were prepared based on the procedure described in ^48^, We previously found that when the high-affinity ACAT inhibitor such as F12511 or K604 are added to cells, the ACAT enzyme stay inhibited by F12511 or by K604, the inhibition does not become diluted by washing the cells, or by the cell homogenization preparation process ^24, 48^. This finding implicates that once the inhibitor binds to the ACAT enzyme, it stays firmly bound for a few hours before dissociating from the enzyme. Based on this finding, we treated animals with the inhibitor by IV injection, perfused the mice after animals were treated (to remove blood contamination from tissues), then prepared tissue homogenates, and monitored ACAT enzyme activity in these tissues, by using the mixed micelles enzyme assay described previously ^24, 39, 52, 56^. (The mixed micelles assay measures the ACAT enzyme activity independent of endogenous lipid composition because it reconstitutes ACAT protein in mixed micelles with defined lipid composition). We injected nanoliposome with F12511 (nanoparticle F) at 5.8 mg/kg, or with K604 at 40 mg/kg, by IV. 4 h after IV, we sacrificed the mice and compared ACAT enzyme activity remaining in various tissues in nanoparticle F or K604 treated mice vs that of the untreated mice. The results (**Fig 1**) showed that F12511 inhibited the ACAT enzyme in the brain, as well as in the adrenal glands, while K604 only inhibited ACAT enzyme in the adrenal glands but not in the brain. This result shows that as part of nanoparticles, F12511, but not K604, is permeable to the mouse brain. This result is consistent with the work of Shibuya et al ^47^ who showed K604, a quite hydrophilic molecule, is impermeable to the brain. It also shows that, even at relatively low dose (5.8 mg/kg), 8 h after IV injection, F12511 is still able to inhibit ACAT activity in the brain by more than 50%.

**Fig 1.**
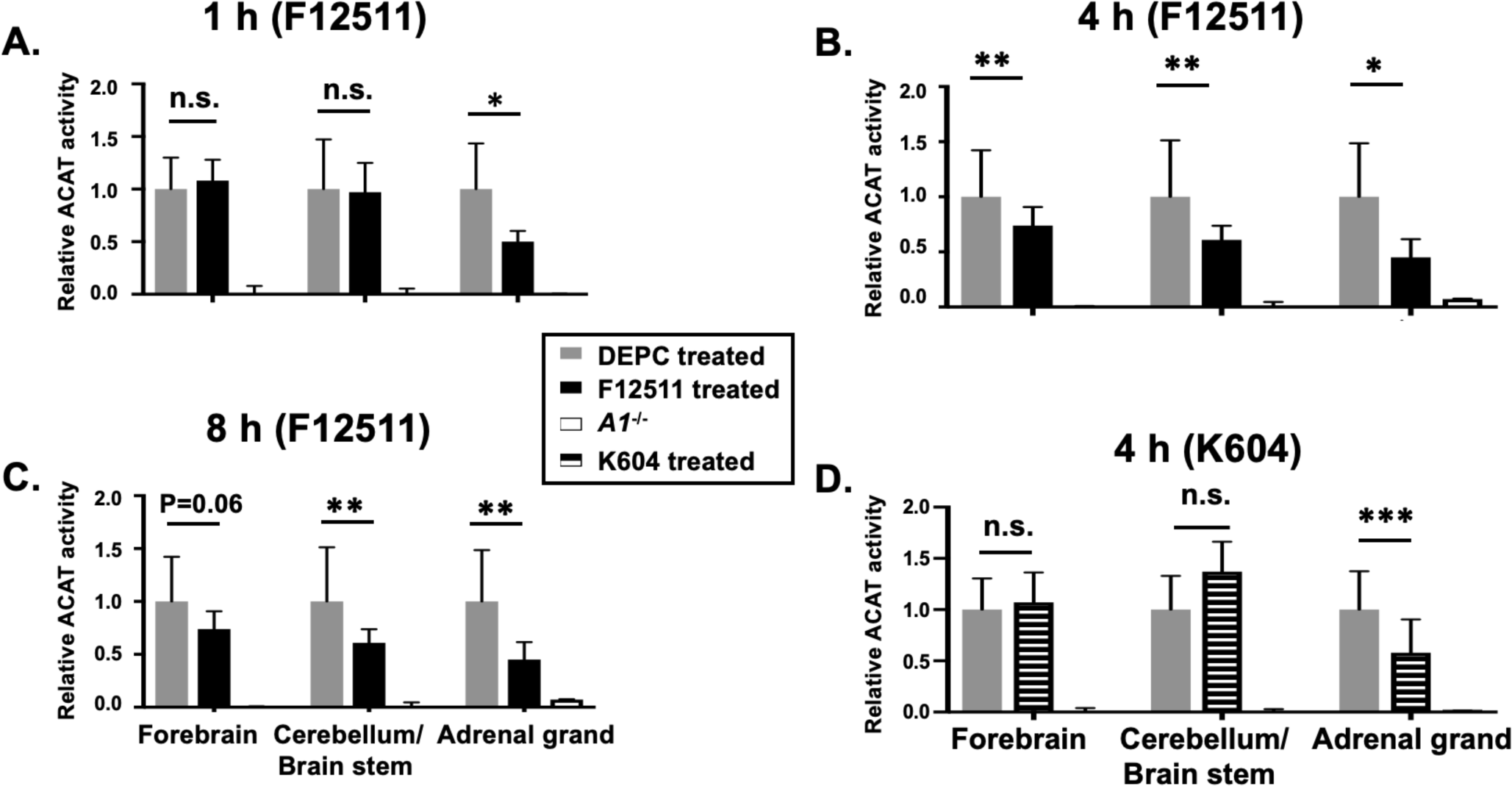
IV injection of nanoparticle F12511 reduces ACAT activities in both adrenal glands and brain, while nanoparticle K604 only reduces ACAT activity in adrenal glands but not in brain. WT mice were IV injected with either nanoparticle F12511 with F12511 at low concentration (30 mM DSPE-PEG2000 with 5 mol% F; 5.8 mg/kg) or with empty nanoparticle at zero time and sacrificed after (A) 1 h, (B) 4 h, (C) 8 h. (D) WT mice were IV injected with nanoparticle K604, with K604 at high concentration (30 mM DSPE-PEG2000 with 40 mol %K) or with empty nanoparticle at zero time and sacrificed after 4 hr. A1−/− refers to Acat1−/− mouse as a negative control. Relative ACAT activity was determined by using the mixed liposome enzyme assay as described in the text and in PMID: 9857049. WT mice at 4 per group were age (4–5 months) and gender matched. N = 1 for A1−/− mouse. n.s., not significant.

To examine if IV injection of nanoparticle F causes preferential accumulation of F12511 in adrenals or in any other tissues, we conducted mass analyses of nanoparticle F by using a LC/MS/MS based procedure described in ^57^. The results (**Fig 2; top**) showed that, after a single IV injection of nanoparticle F at high dose (46 mg F12511/kg), at 4 h, F12511 content in the plasma reached values close to 30 µM. To compare this value with value in the literature: It was previously known that F12511 can be solubilized at high concentration by cyclodextrin. When mice were orally fed with F12511/cyclodextrin complex, the maximal blood F12511 concentration can reach was only 0.75 µM (data retrieved from the Pierre Fabre patent US patent 5990173); i.e., a value less than 30-fold than the method described here can achieve. Results in **Fig 2** bottom graph showed that after a single IV of nanoparticle F, the F12511 content decreased rapidly in the plasma, adrenals, livers, and brains with time (from 4 to 24 h); these results showed by mass spectrometry analyses indicated that there was no evidence for preferential accumulation of F12511 in the adrenals. We also monitored the ACAT enzyme activities in various tissues, 4 to 48 h after a single IV injection of nanoparticle F at 46 mg/kg. The result (**Fig 3**) showed that, similar to the F12511 tissue content analyses results (**Fig 2, bottom**), the ACAT enzyme activity in various tissues (including the adrenal glands) steadily returned to near-normal levels within 24 to 48 h, demonstrating there is no evidence to suggest prolonged, preferential accumulation of F12511 in the adrenals.

**Fig 2.**
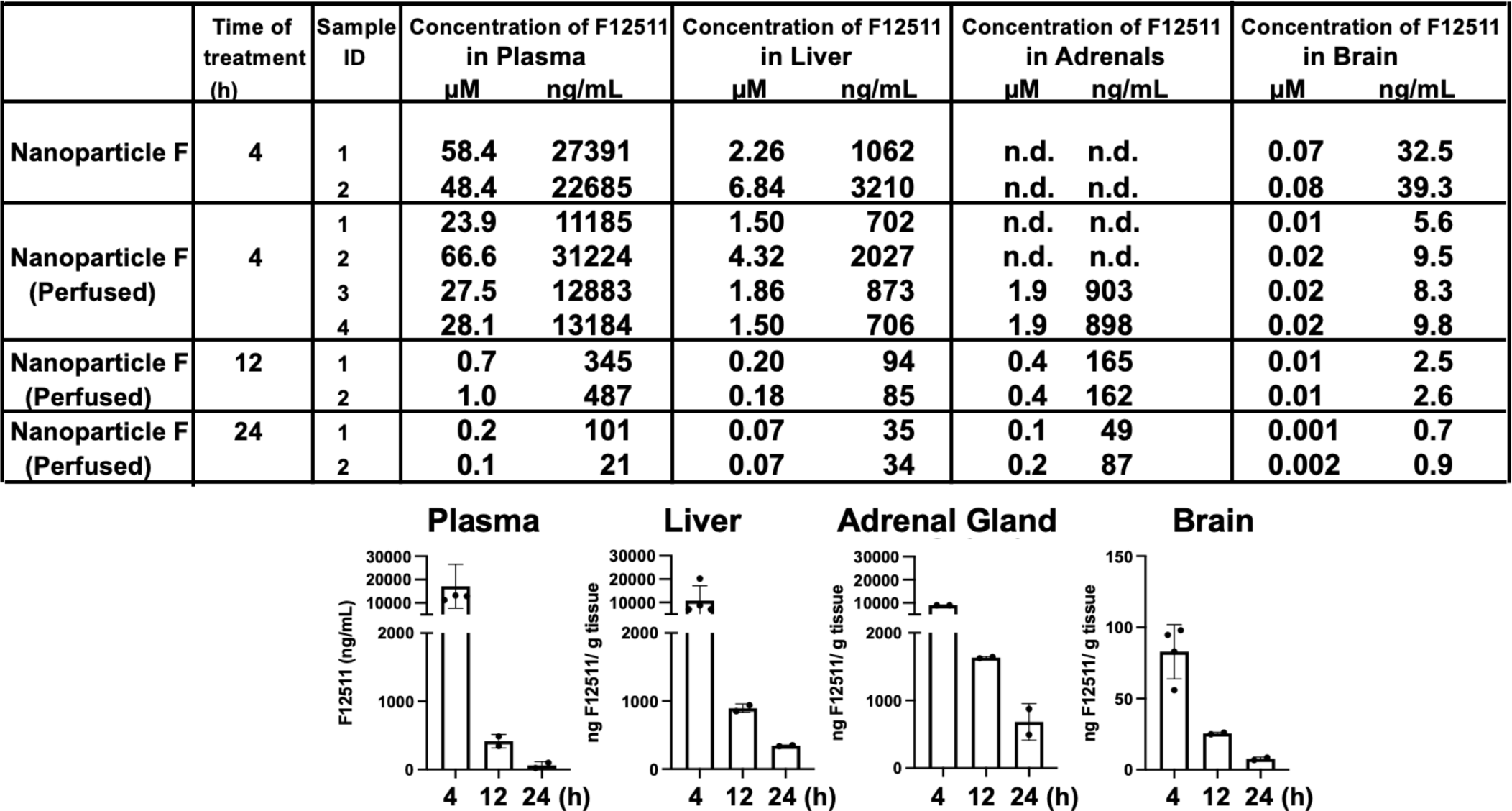
HPLC/MS/MS analyses of F12511. HPLC/MS/MS results (at top panel) shown F12511 content in the mouse plasma, liver, adrenal glands, and brain, after 4, 12, and 24 h post IV delivery of nanoparticle F12511. Most of the mice were with perfusion with PBS for 15 min, to avoid blood contamination of tissues, before sample collections. For each mouse tissue measured, each point represents the average of 3 replicates, with 3 HPLC column injections per replicate. DSPE-PEG_2000_/PC treated mice were also measured by HPLC/MS/MS. As expected, values for F12511 were under the detectable limit. Results from the table on top were replotted in graphs shown at bottom, using results from perfused tissue.

**Fig 3.**
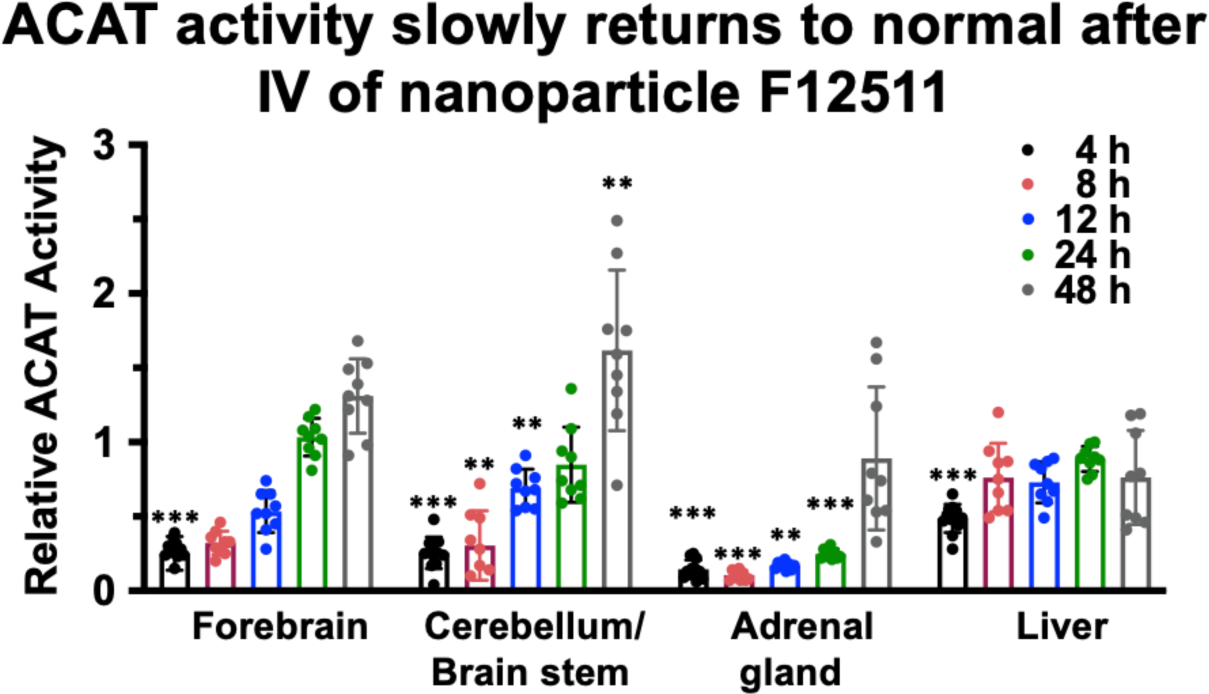
The ACAT activity in forebrain, cerebellum, liver, adrenals gradually return to normal 24 to 48 h after a single IV of nanoparticle F12511. Adult C57BL/6 mice were given single IV of nanoparticle F12511 at high dose (∼46 mg/kg). At various time point indicated, mice were perfused with PBS. The forebrain, cerebellum/brain stem, adrenal gland, and liver were isolated and homogenized to measure ACAT enzyme activity by procedures as described in Fig 1.

### 3.2 IV injection of nanoparticle F does not produce overt systemic or neuro-toxicities *in vivo*

We had previously shown that treating nanoparticle F at very high concentrations to mouse primary neurons in culture did not produce obvious toxicities ^48^. Previous work by others showed that when fed to mice for seven days, certain toxic ACAT inhibitors, such as AZD3988 caused significant reduction in fine vacuolation in cortical regions ^58^. Here, we tested if nanoparticle F would produce toxicity in various tissues by giving IV injections of nanoparticle F to mice at 46 mg/kg once per day for 7 days. Two days after the last injection, we sacrificed the mice and examined the morphologies of various brain regions, the liver, and the adrenal cortices after histochemical staining (**Fig 4**). As shown in **Fig 4 A, B**, no detectable morphological alteration was found in the CNS tissues, or in the liver; nanoparticle with or without F might have induced certain small but detectable vacuolization (**Fig 4C**). The exact cause for this observation is unknown. Adrenal cortical vacuolization in rats and mice ^59^ is characterized by the accumulation of clear vacuoles within cells, mainly in the zona fasciculata but also in the zonae reticularis and glomerulosa. The vacuolization can be focal or diffuse. We speculate the vacuoles seen (**Fig 4C**) may represent accumulations of cholesterol and/or other lipids.

**Fig 4.**
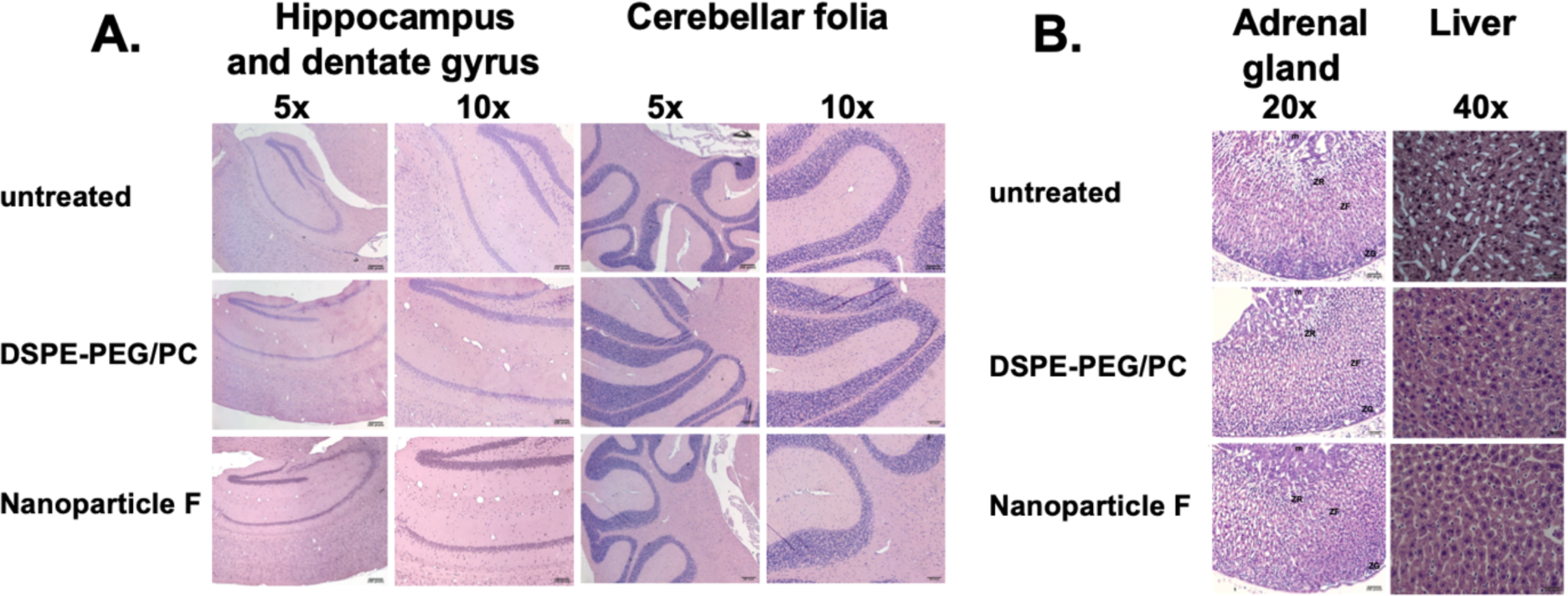
Treating normal mice with nanoparticle F12511 or nanoparticle alone produces no overt morphological alterations in central nervous and peripheral tissues. Representative images from wildtype C57BL/6 mice (n=3-4/group) either untreated or treated by IV injections at once per day with DSPE-PEG_2000_/PC or with nanoparticle F12511 for 7 days. Tissues were collected 48 h after the last injection. **(A)** Images show hippocampus and dentate gyrus regions (left panels). Images on the right panels show cerebellar folia region. **(B)** Images show adrenal glands (left panels), and livers (right panels). For adrenal gland images, symbols used: m=medulla; ZR=zona reticularis; ZF=zona fasciculata; ZG=zona glomerulosa.

### 3.3 Nanoparticle F are present and detectable in the brain by using the fluorescent dye DiR

To monitor biodistribution of nanoparticle, we adapted the procedure from Bishnoi et. al ^55^ and Meng et al. ^60^, by encapsulating the highly fluorescent dye 1,1′-dioctadecyl-3,3,3′,3′-tetramethylindotricarbocyanine iodide (DiR), which is a hydrophobic long-chain dialkylcarbocyanine ^60^. DiR is a lipophilic NIR dye that can be incorporated into lipid bilayer of nanoparticles. ^60, 61^ Moreover, unencapsulated free DiR exhibits only weak fluorescence signal, which does not interfere with signal from encapsulated DiR nanoparticle ^60^. After IV delivery, we isolated different brain tissues including forebrains and cerebellum to produce tissue homogenates and quantitated DiR signal present in these homogenates. The result showed that after 4 h of IV injection, DSPE-PEG/PC/DiR nanoparticle were present in various regions of the brain (**Fig 5**). Quantitation of the results shown in (**Fig 5**) suggested about 0.3-0.5% of nanoparticle injected into the blood entered the brain interior. This value is consistent with the literature reported value ^62^. However, it can only be considered as semi-quantitative because the values reported in **Fig 5** is influenced by tissue environment, which affect the DIR fluorescence signal. To compare the HPLC/MS/MS analyses of F12511, the result showed at 12 to 24 h time point, about 0.5% to 1% of F12511 in the plasma is found in the brain interior (**Fig 2, Table on top**). Together, we speculate that, soon after IV injection, a portion of F12511 may dissociate from the nanoparticle and undergoes rapid degradation by liver, while other portions of F12511 might enter the brain along with the nanoparticle.

**Fig 5.**
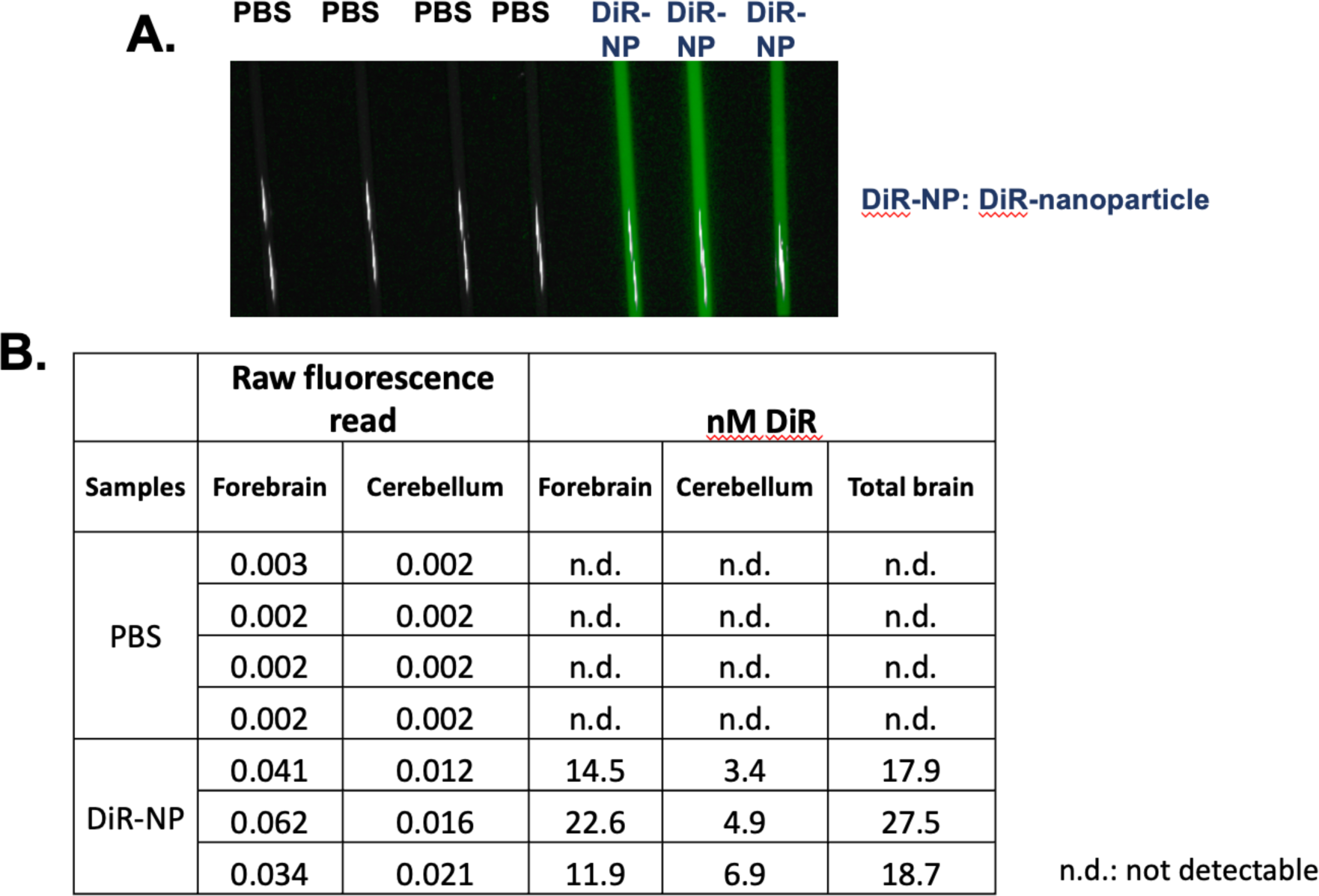
Biodistribution of nanoparticles. 3-month-old mice were injected with 8.3 mM of 1,1’-Dioctadecyl-3,3,3’,3’-Tetramethylindotricarbocyanine Iodide (DIR) encapsulated in DSPE-PEG/PC nanoparticles (NP). Mice were sacrificed after 4 h and perfused with 20 mL of cold 1xPBS. Brains were collected and separated into 2 hemispheres. The right hemisphere was homogenized in lysis buffer and load into capillary tubes for imaging capture on the Pearl 500. **(A)** Representative of mouse brain homogenates in capillary tubes, **(B)** Quantitative analysis of DiR in mouse brain homogenate. All images were analyzed in Fiji. N = 3-4 mice/treatment group. Scale bar = 2500 μm. n.d.: not detectable. Procedure adapted from PMID: 29920064 & PMID: 34514233

### 3.4 Nanoparticle F suppressed amyloid-beta (Aβ) and diminished the levels of both unphosphorylated and hyperphosphorylated Tau (HPTau)

We had previously shown that *Acat1/Soat1* gene knock out (KO) in the 3xTg AD mouse model ^50^ reduced mutant full length hAPP, suppressed the level of Aβ (Bryleva, Rogers et al. 2010), and diminished the levels of unphosphorylated mutant human Tau, but not hyperphosphorylated mutant human Tau (HPTau) ^32^. Here we tested the therapeutic efficacy of nanoparticle F in the 3xTg AD mouse model. In this model, Aβ, tau phosphorylation, and neuroinflammation become pronounced between 12–20 months of age ^63^. We tested nanoparticle F in mice with advanced-age (16–20 months). Male and female mice were left untreated or were treated with daily IV injections (alternate between tail vein and retro-orbital; 200 µL per injection) of nanoparticle F (∼46 mg/kg F12511), or DSPE-PEG_2000_/PC (nanoparticle alone) for 2 weeks. After treatment, mouse tissues were collected and snap-frozen until ready for tissue homogenization, western blot analyses, ELISA and Luminex assay. The results showed that nanoparticle F significantly reduced mutant hAPP as determined by western blot (**Fig 6A**) and reduced total acid extractable Aβ1-42 as determined by ELISA assay (**Fig 6B**). Interestingly and unexpectedly, nanoparticle alone also reduced full-length hAPP, although the effect is milder than that of nanoparticle with F12511 (**Fig 6A**), suggesting the nanoparticle without F12511 may exert a certain suppressive effect on full-length hAPP. We noted that, in **Fig 6**, there was a large variability in the expression levels of mutant hAPP, and Aβ1-42 levels among the nanoparticle-alone/vehicle treated and the untreated groups. We suspect that the variability might be caused by the variability in ages of these mice used (16–20 months of age).

**Fig 6.**
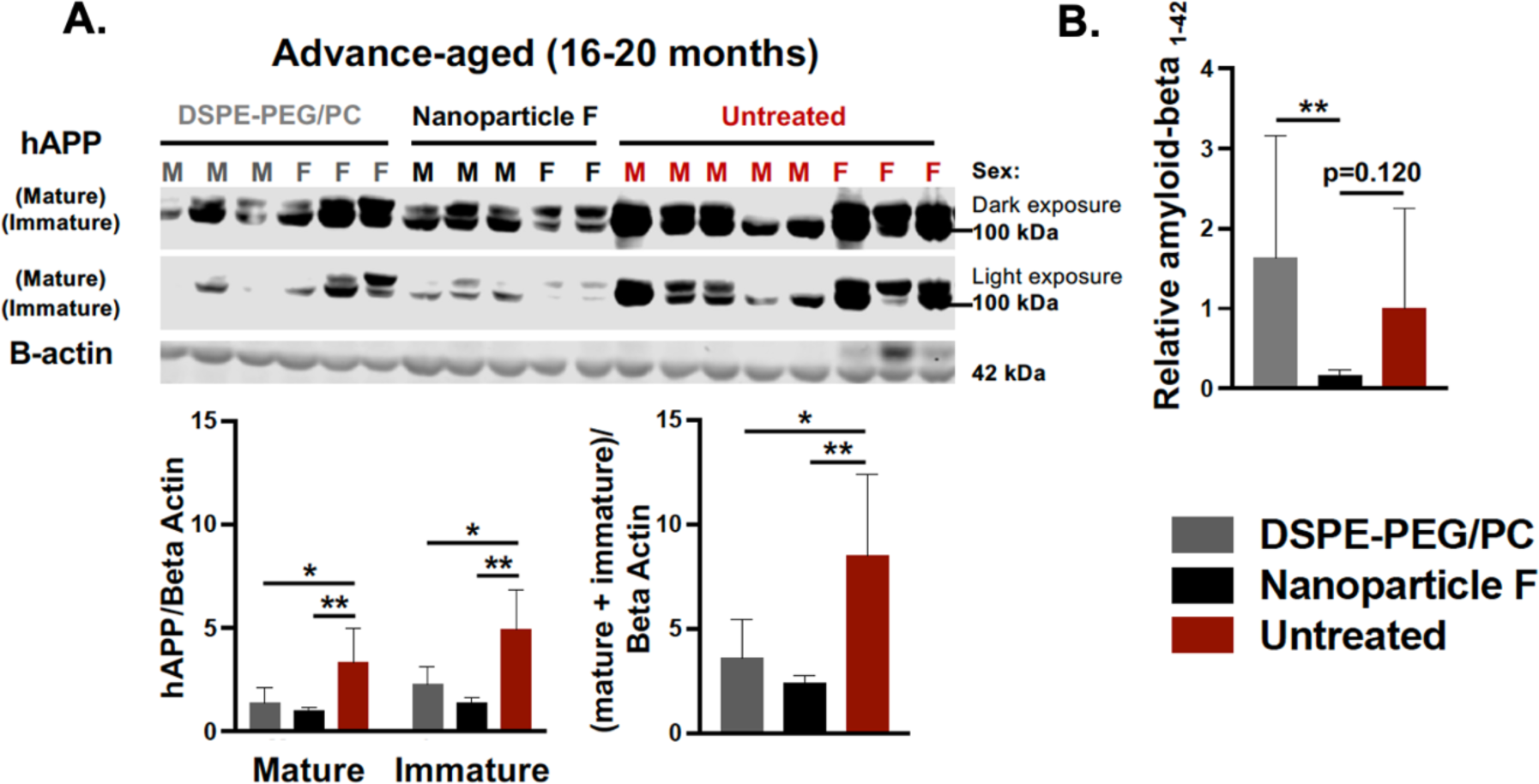
IV injections of nanoparticle F12511 reduce mutant hAPP and reduce total acid extractable Aβ1-42 in advance-aged 3xTg AD mice. Male (M) and female (F) 3xTg AD mice at 16-20 months of age were either untreated or treated with DSPE-PEG_2000_/PC (vehicle) or with nanoparticle F (F12511 at 46 mg/kg) as indicated with daily IV injections once daily for 2 weeks. Mouse brain homogenates were prepared according to procedure described in PMID: 15980612 and PMID: 20133765. (**A**) Western blots to monitor the immature and mature forms of hAPP (two adjacent bands at 105-kDa and 115-kDa; PMID: 20133765), using mouse monoclonal antibody anti-6e10 (recognizing amyloid Aβ; 1:5000, from Covance). Results are shown as light and dark exposures. Two-way ANOVA was conducted to analyze the signals for mature and immature bands individually. One-way ANOVA was used to analyze total signals comprising of both mature and immature bands. Mouse anti-beta actin antibodies were used as loading controls. (**B**) ELISA assay using mouse monoclonal antibody anti-6e10 to monitor total formic acid extractable Aβ1-42 levels in brain homogenates. Procedure for Aβ1-42 extraction was described in PMID: 15980612 and PMID: 20133765. ELISA plates were from Invitrogen. Each sample was measured in quadruplicate. Results are presented as Aβ1-42 signal intensity from mice treated with DSPE-PEG_2000_/PC (nanoparticle/vehicle) or with nanoparticle F relative to that of untreated control using one-way ANOVA analysis. N.S. not significant; p<0.001 ***, p<0.01 **, p<0.05*.

Besides amyloid deposition, 3xTg AD mice also develop tau pathologies. We measured changes in tau by performing western blots using two antibodies: HT7 (which recognizes total human tau; hTau), and AT8 (which recognizes human tau phosphorylated at Ser202/Thr205). The results show that, when compared with untreated mice, only nanoparticle F, but not nanoparticle alone, significantly reduced total human tau levels (**Fig 7A**). Interestingly and unexpectedly, both nanoparticle F and nanoparticle alone significantly reduced phosphorylated tau HPTau levels (**Fig 7B**). These results again suggest that the nanoparticle alone may exert a certain suppressive effect on HPTau level.

**Fig 7.**
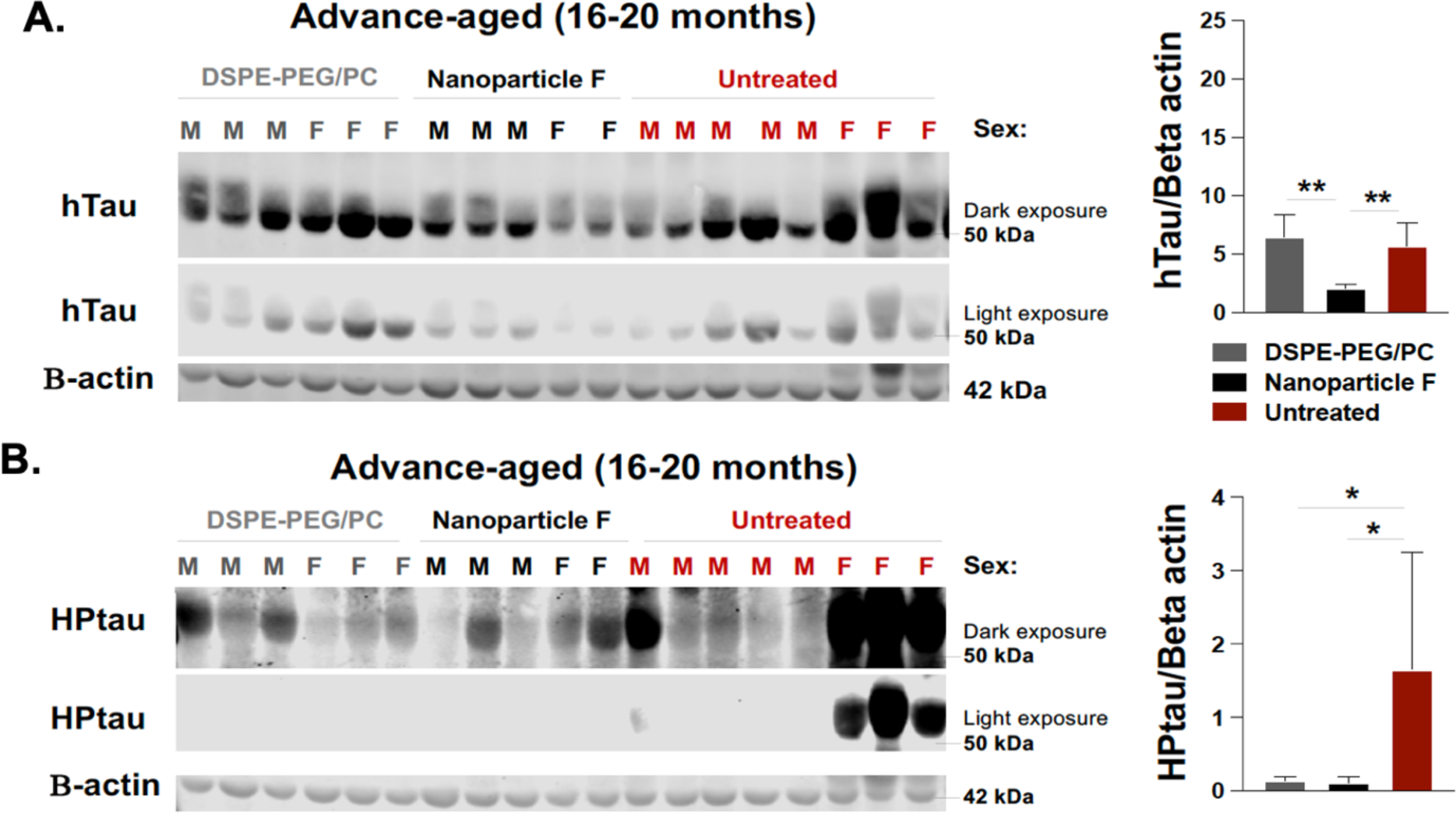
Nanoparticle F12511 reduces total unphosphorylated mutant human tau (hTau) and hyperphosphorylated human tau (HPTau), while nanoparticle alone reduces HPTau but does not reduce htau in advance-aged 3xTg AD mice. The same brain homogenate samples described in Fig 6 were used for Western blot analyses to monitor hTau (A) and HPTau (B), according to procedure described in PMID: 25930235. Mouse anti-HT7 for total unphosphorylated human tau (∼50 kDa) and mouse anti-AT8 for hyperphosphorylated human tau (∼55-60 kDa) were from Thermo Fisher Scientific; mouse anti-beta actin (42 kDa) antibodies served as loading control. Dark and light exposure of each blot is shown. One-way ANOVA was conducted for statistics. N.S. not significant; *p*<0.001 ***, *p*<0.01 **, *p*<0.05 *.

### 3.5 Nanoparticle F attenuate neuroinflammation in the aging 3xTg mice

The aging 3xTg AD mice exhibit chronic neuroinflammation ^63^. Here we tested if nanoparticle F may attenuate neuroinflammation in these mice at 16-20 months of age. After a two-week IV injection to these mice, we prepared whole brain homogenates and analyzed 31 different mouse cytokines using Milliplex® technology ^64^. The result in **Fig 8A** compares the average cytokine expression level between different experimental treatment group. Our data suggested that, when compared to nanoparticle treatment alone, treatment with nanoparticle F showed a decrease in trend of 27 out of 31 analyzed cytokines, of which mostly are pro-inflammatory in nature **(first two column from the left, Fig 8A)**. In some cytokines, such as TNF-α and Eotaxin, nanoparticle and F12511 work additively to reduce their levels. (**Fig 8A**) To further examine the effect of nanoparticle and nanoparticle F in the aging 3xTg AD mice, we plot individual cytokines data that were previously shown in literature (as reviewed in ^65–67^ to be related to neuroinflammation and Alzheimer’s disease including: IL-1α, IL-1β, IL-6, TNF-α, IL12-p40, Eotaxin, MIP-1α, MCP1 (**Fig 8B**). For cytokines such as IL-1α, IL-1β, IL-6, IL-12p40, MIP-1α, MCP1, treatment with nanoparticle F reduces their level compared to treatment with nanoparticle alone, where minimal effect from nanoparticle alone was observed when compared to untreated 3xTg AD mice (**Fig 8B**). In cytokines such as TNF-α and Eotaxin, nanoparticle alone treatment reduced cytokines level compared to untreated 3xTg AD mice (**Fig 8B**). Treatment with nanoparticle F further reduces cytokines level compared to nanoparticle alone treatment, highlighting the synergetic effect of nanoparticle and F12511 (**Fig 8B**). We used Volcano plots to illustrate the difference between all 31 cytokines expression between each treatment group (shown in **Supplementary Figure 1**). We note that the effects of nanoparticle and nanoparticle F on cytokines expression are cytokine specific. Together, these results suggested that treatment with nanoparticle alone and nanoparticle F reduces most pro-inflammatory cytokines expression in the aging 3xTg AD mice, human Tau and HPTau were previously shown to be highly inflammatory in the brain and removal of these aggregated protein species could restore proper brain function through ameliorating neuroinflammation. We suspect that the effect of nanoparticle and nanoparticle F treatment on inflammatory cytokines could be in part, due to the ability of nanoparticle and nanoparticle F treatment in reducing Aβ, human Tau and HPTau level.

**Figure 8.**
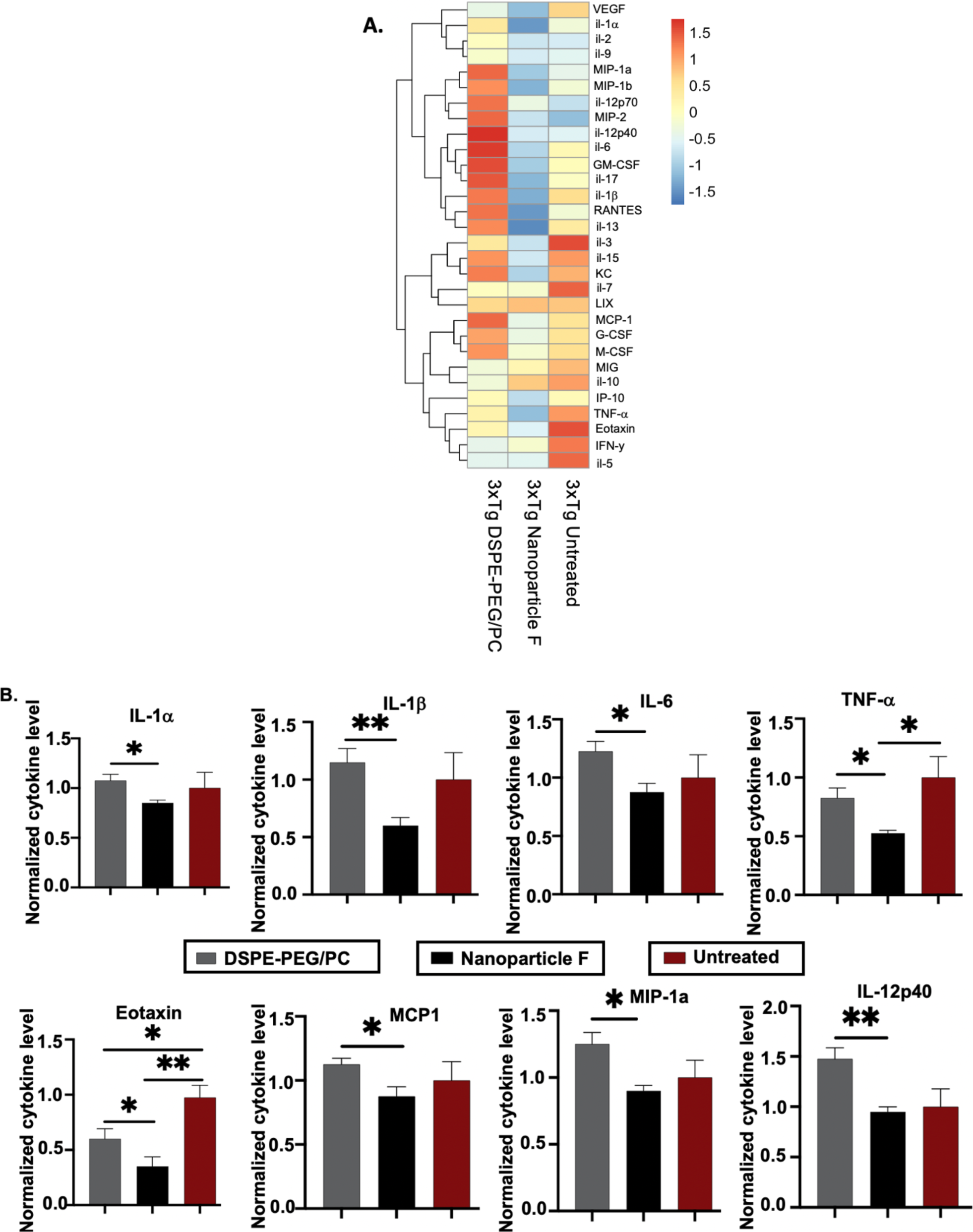
DSPE-PEG/PC and nanoparticle F alter pro-inflammatory cytokines profile in 3xTg AD mice (16-20 month; after 2 weeks daily IV/RO injections. Mice were perfused with 20 mL of cold 1xPBS, brains were collected and homogenized. Forebrain homogenates were analyzed using MILLIPLEX MAP Mouse Cytokine/Chemokine Magnetic Bead Panel (32-plex) and data were normalized using total protein content determined by Lowry assay. **(A)** Heatmap visualizing average cytokines readings from several biological replicates of each treatment group with a Z-score transformation. **(B)** Alzheimer’s related cytokines were plotted individually with 3xTg mice forebrain cytokines content. For B, values obtained from each individual cytokine from PBS injected animal is normalized to 1. Unpaired student t-test (two-tailed) were performed, \**P* < 0.05, \*\**P < 0.01* n = 4 mice/group.

## 4. Discussion

In this study we aimed to characterize nanoparticle F treatment *in vivo* by pharmacokinetic studies using ACAT enzymatic activity assays and HPLC/MS/MS analyses. We provided evidence that nanoparticle F is able to permeate the blood brain barrier (BBB). We also showed that by a single intravenous (IV) injection, nanoparticle F at ∼46 mg/kg concentration inhibited ACAT in the brain by ∼70% for up to 12 h, and in the periphery for up to 24 h. Importantly, ACAT activity was restored fully in all tissues examined at 48 h after injection, showing that F12511, a hydrophobic compound, was not accumulating in the tissues that we had examined. Previously, F12511 passed the clinically safety test and did not cause adrenal toxicity in animals ^36^. To confirm that nanoparticle F does not cause overt toxicity, we treated mice by daily IV for 7 days with nanoparticles, with or without F12511. Afterwards, histological examinations in the adrenal glands, liver, and brain tissues were performed, the results showed no detectable major morphology changes or tissue damages occurred.

Once nanoparticle F was characterized in wildtype mice, therapeutic efficacy for AD treatment was investigated. The results showed nanoparticle F treatment, as well as control nanoparticles, provided benefits in advanced-age 3xTg AD mice. The rather unexpected and exciting finding from this work is that the nanoparticle alone, and F12511 may produce an additive effect on amyloidopathy and on tauopathy: Based on western blot results, nanoparticle F and vehicle nanoparticles both reduced full-length mutant hAPP and phosphorylated human tau; the effect was strongest with nanoparticle F. Furthermore, only nanoparticle F, but not nanoparticle alone, reduced Aβ1-42 and total mutant human tau. Alzheimer’s pathology drives neuroinflammation through pro-inflammatory cytokines expression in the brain. (Reviewed in ^65^) Treatment with nanoparticle F significantly reduced classical pro-neuroinflammatory cytokines in 3xTg AD mice brains such as IL-1β, IL-6 and IL-12p40, as well as other pro-inflammatory cytokines comparing to nanoparticle alone treatment. In cytokines such as TNF-α and Eotaxin, treatment with nanoparticle and nanoparticle F produce additive effect to further dampen proinflammatory cytokines level in 3xTg AD mice brains. Our cytokines analyses result from 3xTg AD mouse brains further corroborate the effects observed on reducing full-length mutant hAPP, Aβ1-42, unphosphorylated and phosphorylated human tau, supporting that nanoparticle alone and F12511 produce additive effects. How do the additive effect occur mechanistically requires further investigation.

Overall, our results show that the DSPE-PEG_2000_ and PC not only increased the amount of F12511 that was encapsulated into the nanoparticles, but they also produced benefit in therapeutic treatment. DSPE-PEG and PC have been previously suggested to have some benefit in certain disease model systems. Notably, Brown and colleagues tested the effects of using DSPE-PEG micelles in combination with cyclodextrin as a potential therapy for Niemann Pick Type C (NPC) disease, an adolescent lysosomal storage disorder ^68^ The use of 2-Hydroxy-propyl-b-cyclodextrin (HPbCD) has been used in clinic for treatment of NPC-Type 1 but required multiple high doses, which resulted in ototoxicity or hearing loss. Utilizing DSPE-PEG not only had an effect itself but worked synergistically with HPbCD in cell culture ^68^. A separate study looking at AD mouse model utilized liposomes with phosphatidic acid (PA), which is a metabolite of PC. PA was shown to interact with Aβ1-42 ^69^, and was utilized, along with transferrin and a neuroprotective peptide, to generate liposomes to reduce amyloid in AD mouse models ^70^. Yang and colleagues used a trans well migration assay, where microglia can migrate to different chambers. The results showed that microglial chemotaxis towards the Aβ-oligomer wells was increased in both PA liposomes with and without the neuroprotective peptide of interest, suggesting the PA liposome itself was beneficial. These PA liposomes reduced amyloid; however, to our knowledge, the effect of liposomes has not been documented with tauopathy.

One important note is the possibility for the nanoparticle F to exhibit more pronounced effect in the advanced-age range, due to BBB integrity. It has been well established that BBB integrity and function are compromised in certain diseases, including neurodegenerative diseases like AD. In addition, in aged mice there is a dysfunction of BBB tight junctions, and this is accompanied by an increase in neuroinflammation ^71^. We speculate that the benefits seen in the advanced-age group may be due to a “leaky” BBB that allows more nanoparticle F to enter the brain parenchyma. As mentioned above, F12511 was detectable in the brain by HPLC/MS/MS at a low concentration (∼10-20 nM); however, this value was obtained using normal adult mice at 2-4 months of age. With a compromised BBB, it is possible that the F12511 concentration in the brain is a lot higher. Future studies are needed to test if modifying the chemical structure of F12511 to increase its BBB permeability.

We had previously shown that, in the lipopolysaccharide (LPS) induced acute neuroinflammation mouse model, genetic KO of ACAT1 in myeloid cell lineage attenuated neuroinflammation induced by LPS ^20^. Mechanistic studies showed that ACAT1 blockade act in part by increasing the endocytosis of the receptor TLR4 that mediate LPS mediated proinflammatory signaling cascade at the plasma membrane ^20^. Acute neuroinflammation and chronic inflammation share many causal factors in common ^72, 73^. We speculate that nanoparticle F produce anti-inflammation in 3xTg AD mice may occur by suppressing Aβ and/ or Tau / hyperphosphorylated Tau, which are known to cause chronic proinflammatory responses. Additionally, it may also act on alter the fate of TLR4 in microglia or astrocytes. Currently, we do not know how nanoparticle alone act *in vitro* and *in vivo*.

Fisher and colleagues showed that in monocyte/macrophages, acute treatment of apoA1 *in vitro* and *in vivo* produce anti-inflammatory responses by promoting cholesterol efflux at cholesterol-rich lipid rafts, independent of the lipid efflux protein ABCA1 ^74^. We suspect the action of nanoparticle alone may act in similar manner as apoA1: by promoting cholesterol efflux at the lipid rafts region of plasma membrane.^20, 75^ We speculate that blocking ACAT1 may increase cholesterol content at the lipid raft domain in various membrane organelles, and it maybe through the cholesterol rich lipid raft domain that F12511 and nanoparticle alone act synergistically to benefit AD and other neurodegenerative diseases. Future investigation will be required to dissect the mechanism(s) involved in nanoparticle F in suppressing Aβ and/ or Tau / hyperphosphorylated Tau, and in neuroinflammation.

## Author contributions

ADLT, TH, KSS, CCYC, TYC designed research; ADLT, TH, WFH, DN, DCP, SP, YC performed research; ADLT, TH, CCYC, LL, WFH analyzed data; TYC, CCYC, ADLT and TH wrote and edit the manuscript.

## Acknowledgements

We would like to thank the following funding sources: National Institute of Neurological Disorders and Stroke (NINDS) F31NS110317-02 (ALD); National Institute on Aging (NIA) R01AG063544 (TYC and CC); Dana’s Angels Research Trust (to TYC and CC), and NIH IDeA award for Dartmouth BioMT (P20-GM113132), Dartmouth PhD Innovation Program (to TNH), Dartmouth Cancer Center Clinical Pharmacology Shared Resources CCSG 5P30CA021308. Brain Luminex experiment were carried out in DartLab, the Immune Monitoring and Flow Cytometry Shared Resource at the Norris Cotton Cancer Center at Dartmouth, with NCI Cancer Center Support Grant 5P30 CA023108-41.

We would like to thank Hermes Yeh for the suggestion of testing perfusion vs non-perfusion methods, thank the Center for Comparative Medicine and Research at Dartmouth Specifically, we would like to thank Kirk Mauer, Eric DuFour and Catherine Bennett, for their help and training in all of the injection methods used in this research. We would like to thank Junghoon Lee for critical reading of this manuscript.

## Ethics approval and consent to participate

All the experiments were performed in accordance with legal and institutional guidelines and were carried out under ethics, consent and permissions of the Ethical Committee of Care and Use of Laboratory Animals at Geisel School of Medicine at Dartmouth.

## Consent for publication

Not applicable.

## Availability of data and materials

The authors declares that the relevant data are included in the article.

---

**Supplementary Figure 1.**
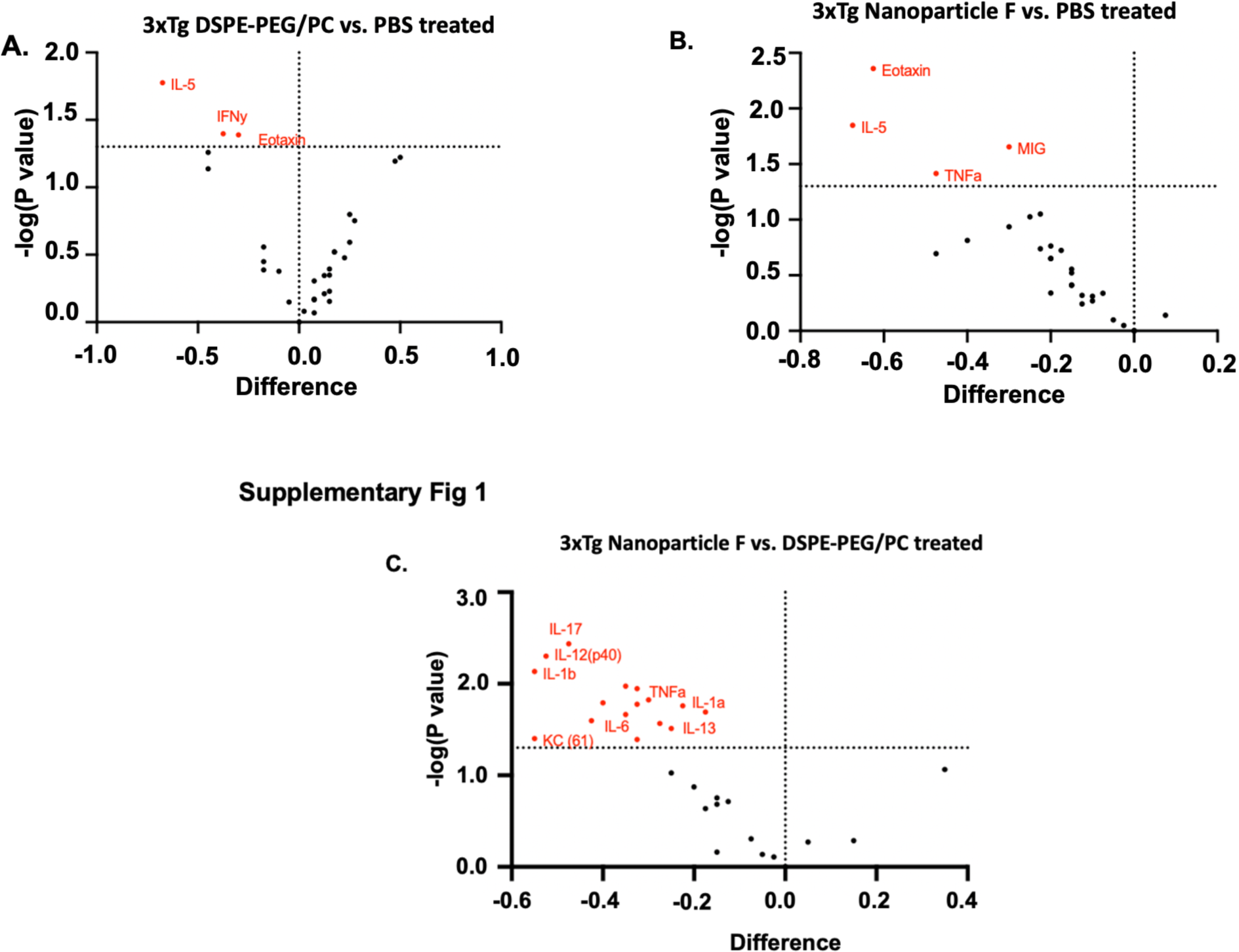
Brain cytokines differentiate upon treatment with DSPE-PEG/PC and nanoparticle F. Volcano plot showing cytokines change upon **(A)** DSPE-PEG/PC treatment vs. PBS, **(B)** nanoparticle F vs. PBS, **(C)** nanoparticle F vs. DSPE-PEG/PC in 3xTg mice. Horizontal dashed line is determined at p = 0.05. Data points above the dash line are statically different at p < 0.05 determined by unpaired student t-test. n = 4 mice/group.

